# Conformational Transition Pathways in Major Facilitator Superfamily Transporters

**DOI:** 10.1101/708289

**Authors:** Dylan Ogden, Kalyan Immadisetty, Stephanie Sauve, Mahmoud Moradi

## Abstract

The major facilitator superfamily (MFS) of transporters contains three classes of membrane transporters: symporters, uniporters, and antiporters. Despite utilizing a variety of transport methods, MFS transporters are believed to undergo similar conformational changes within their distinct transport cycles. Although the similarities regarding conformational changes between the classes of MFS transporters are noteworthy, the differences are also valuable because they may explain the distinct functions of the classes within the MFS. Here, we have performed a variety of equilibrium and non-equilibrium all-atom molecular dynamics (MD) simulations of the bacterial proton-coupled oligopeptide transporter (GkPOT) and the human glucose transporter 1 (GluT1). To compare the similarities and differences of the conformational dynamics found within the three different classes of transporters we have also referenced previous simulations involving the glycerol-3-phosphate (GlpT) transporter. All of the proteins discussed here were simulated in the *apo* state in explicit membrane environments. Our results suggest a very similar conformational transition for all transporter types involving interbundle salt-bridge formation/disruption coupled with the orientation changes of transmembrane (TM) helices, specifically H1/H7 and H5/H11, resulting in an alternation in the accessibility of water at the cyto- and periplasmic gates.

## Introduction

Membrane transporters facilitate the exchange of materials across lipid bilayers. Among these transporters, the MFS is the largest family of secondary membrane transporters, comprised by more than 10,000 members.^1–10^ Examples of MFS transporters that have been studied structurally include the lactose permease, a sugar-porter from *Escherichia coli*,^11–14^GlpT,^15^ the xylose transporter,^16^ and the multidrug transporter EmrD.^17^

All MFS transporters share a common 12 transmembrane (TM) helical structure, consisting of an N- and a C-bundle domain, displaying a twofold pseudo-symmetry.^4,6,10,18–20^ However, MFS transporters function in a number of different ways including active and passive transport. The latter function is uniport (passive transport of the substrate along its concentration gradient) and the former function can be either symport or antiport^15,18,21^ (active transport of the substrate against its concentration gradient coupled with the transport of the co-transported species along their concentration gradients in the same or opposite direction, respectively). Despite these distinct functions, MFS transporters all share a common overall mode of function known as the “alternating access” mechanism, which is shared with other membrane transporters.^1,10,18,21–29^ According to this mechanism, the binding site is never exposed to both sides of membrane at the same time; instead, the protein alternates between an inward-(IF) and an outward-facing (OF) conformation. A more specific model of alternating access in MFS transporters has been proposed which is known as the “rocker-switch” mechanism.^15,29^ In order for transport to ensue using the rocker-switch mechanism, the N- and C-bundle domains undergo concerted conformational changes that expose the binding site to the two sides of membrane alternatively and provides a mechanism for substrate translocation (Figure 1). In addition to the IF and OF states, the transport mechanism also involves occluded intermediates where the binding site is not accessible to either side of the membrane. Models and structures of other members of the MFS have been solved in an occluded state.^14,30–32^

**Figure 1:**
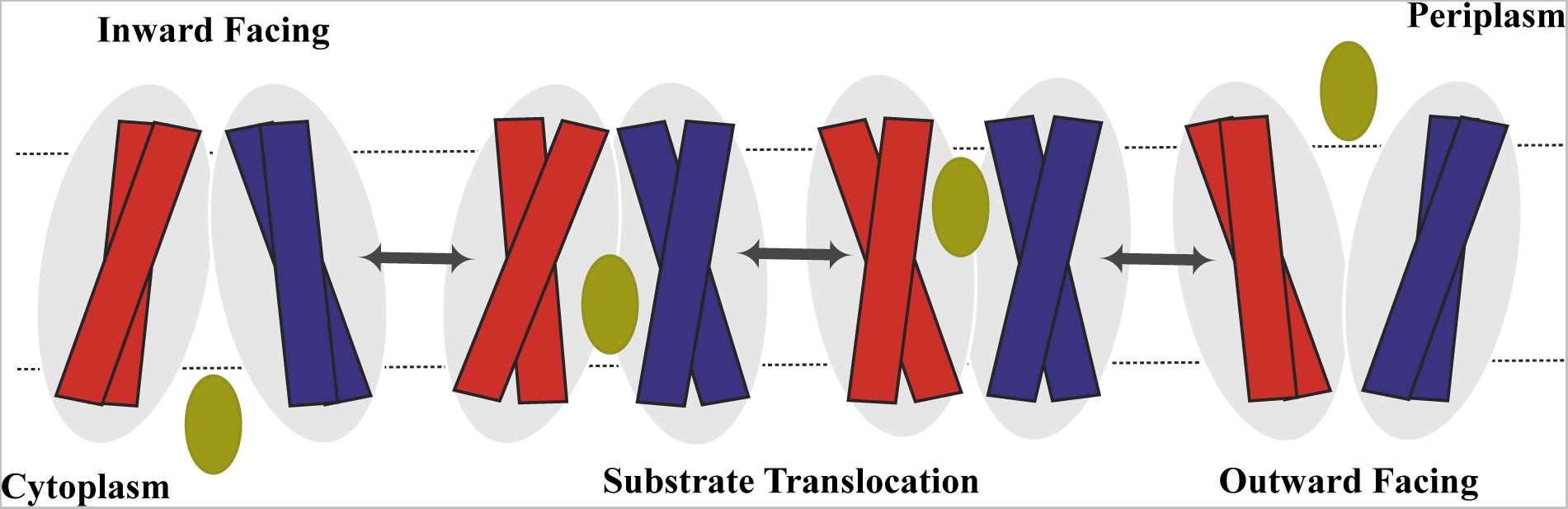
Schematic representation of the MFS facillitative diffusion of a substrate shown in green across a cell membrane using the rocker-switch alternating access mechanism. The N-terminal domain is shown in blue and the C-terminal domain is shown in red.

Structural similarities of the MFS transporters suggest that these transporters may undergo similar conformational changes during their inter-conversions between the IF and OF states; although the coupling of these conformational changes to different binding/unbinding events is likely to be quite different in various symporters, uniporters, and antiporters. In order to study the similarities and differences of the structural changes in the three different classes of MFS transporters, we present a comparative view of IF*↔*OF conformational changes of three proteins from three different classes of MFS transporters, including a bacterial proton-coupled oligopeptide transporter, namely GkPOT,^33,34^ the human GluT1,^35,36^ and the bacterial GlpT.^15^ We specifically limit the discussion to the *apo* state of these proteins; however, we are aware that a more complete picture can be provided if the full transport cycle is simulated with substrates included. This would include the binding, unbinding, and translocation of substrates along with their cotransported species. Nevertheless, the study of *apo* protein IF-OF transition is the first step in characterizing the structural changes of MFS transporters within their transport cycle which is the subject of the current study.

GluT1 is a uniporter belonging to the sugar porter subfamily of MFS transporters that mediates the uptake of glucose by passively transporting it along its concentration gradient.^35^ In GluT1, the N and C domains are connected by an intracellular helical bundle (ICH) which comprises four short *α*-helices. The ICH domain is observed in the structures of other sugar porter members, such as the xylose transporter, but not in other MFS transporters, suggesting that the ICH may be a unique feature of the sugar porter subfamily. The ICH in sugar porter members acts as “latch” to secure closure of the intracellular gate in the OF conformation.^36^ Although GluT1 is a passive transporter, recent structural studies suggest that the transport mechanism follows an alternating access mode of function similar to active membrane transporters.^36–38^

Proton-coupled oligopeptide transporters (POTs) are symporter members of the MFS superfamily. POTs uptake small peptides and peptide like molecules to the cell across the cell membrane using the inward directed proton electrochemical gradient as the source of energy for their active symport function.^39–41^ A very important feature among POTs is substrate promiscuity^42^ that is attributed to the binding site which accommodates a range of peptides and peptide like molecules in multiple orientations. ^43^ Mammalian POTs have yet to be crystallized; however, several bacterial POT members have recently been crystallized including PepT*_so_*,^43,44^ PepT*_st_*,^33,45^ PepT*_so_*_2_,^34^ and GkPOT.^40^ Particularly, the POT transporter from *Geobacillus kaustophilus* bacterium has been crystallized in a lipidic environment to a remarkably high resolution of 1.9 Å.^40^

On the other hand, GlpT is an antiporter member of the MFS superfamily that uptakes glycerol-3-phosphate using the outward directed inorganic phosphate concentration gradient as the source of energy for its active antiport activity.^15,46^ The binding of inorganic phosphate facilitates the IF*→*OF state transition, and the replacement of glycerol-3-phosphate reverts the protein back to the IF conformation. ^4,47^

MD simulations have been a useful tool in the extensive study of proteins including membrane transporters such as GkPOT,^48,49^ GluT1,^50^ and GlpT.^6^ Unfortunately, due to the short timescales of typical unbiased MD simulations, this technique is often incapable of capturing large-scale conformational changes. To be able to sample such conformations, enhanced sampling techniques are typically sought after, due to their ability to capture longer timescales. These include simulations such as steered molecular dynamics (SMD) ^51^ or targeted molecular dynamics (TMD).^52^ Admittedly, the transition between the IF and OF conformations may be sampled using unbiased MD for certain proteins. Recent work has shown a complete transition of GluT1 in 1-1.5 *µ*s.^50^ For slower IF*↔*OF transitions, we have proposed a computational recipe that allows for reconstructing the IF*↔*OF transition of various timescales.^6,53,54^ We have recently improved this protocol by employing a Riemannian description of conformational landscape of protein.^55^ The conformational transition pathways of GlpT has already been reported previously for both the *apo* and phosphate-bound forms.^6^Here we compare the IF*↔*OF conformational transition pathway of *apo* GkPOT and GluT1 to that of GlpT and discuss the similarities and differences of these pathways in detail.

## Theoretical Methods

We have employed all-atom MD to study the IF*↔*OF conformational transitions of MFS transporters in the *apo* state. In each case, a membrane-embedded model of the wild-type *apo* protein in its respective IF state was used based on the available crystal structure of protein (PDB: 4IKV^40^ and 4PYP^36^ for GkPOT and GluT1, respectively).

Unbiased simulations of GkPOT were previously simulated by Immadisetty, et al.^48^ and were used as a starting structure for the biased simulations presented here. Regarding the simulation set up of the unbiased simulations, the protein was placed in 1-Palmitoyl-2-oleoylsn-glycero-3-phosphoethanolamine lipids and all the crystal structure waters were removed. The protocols for generating this model have been previously reported in detail.^48^ Similarly, Moradi et al.^6^ previously simulated GlpT (PDB ID: 1PW4^4^) in 1-Palmitoyl-2-oleoyl-snglycero-3-phosphoethanolamine lipids and all the crystal structure waters were removed. Results of the GlpT simulation is referenced throughout this paper. For GluT1, CHARMM-GUI^56,57^ was used to build the simulation system. The mutated residues in the GluT1 crystal structure (PDB: 4PYP)^36^ were substituted with the wild-type amino acids and GluT1 was placed in a lipid bilayer of 1-Palmitoyl-2-oleoyl-sn-glycero-3-phosphocholine, solvated with TIP3P waters,^58^ ionized with 0.15 M KCl, and placed in a box of *≈*106*×*105*×*104 Å^3^.

We used the CHARMM36m all-atom additive force field to describe all molecules^59,60^ and all MD simulations were performed using NAMD 2.11.^61^ Prior to equilibrium runs, each system was energy minimized for 10,000 steps using the conjugate gradient algorithm^62^ and relaxed using a *∼*1 ns multi-step restraining procedure (CHARMM-GUI’s default procedure for membrane proteins^56^). This initial relaxation was performed in an NVT ensemble while all production runs were performed in an NPT ensemble. Simulations were carried out using a 2 fs timestep at 310 K using a Langevin integrator with a damping coefficient of *γ* =0.5 ps*^−^*^1^. The pressure was maintained at 1 atm using the Nośe-Hoover Langevin piston method.^63,64^ The smoothed cutoff distance for non-bonded interactions was set to 10*−*12 Å, long-range electrostatic interactions were computed with the particle mesh Ewald method, ^65^ and periodic boundary conditions were used in all simulations. Here, we performed two sets (replicas) of equilibrium MD simulations for GluT1 where each replica was equilibrated for 700 ns. The simulation times employed were long enough to capture the IF*→*OF transition in the *apo* GluT1.

For GkPOT, the starting point for the biased simulation presented here was the equilibrated model from an unbiased simulation reported previously by Immadisetty, et al., referred to as Set 1 ^48^ in which GkPOT is in the *apo* state and residue E310 is unprotonated. Using this model as input, we then performed SMD simulations using various biasing protocols, described in detail in Supporting Information. Here we only present the results of the most successful protocol which uses a biasing potential based on the orientation of TM helices H1, H2, H7, and H8 to induce the IF*→*OF transition of *apo* GkPOT. The simulation time for this SMD protocol was 100 ns. During this simulation, a harmonic restraint was imposed on the orientation quaternions of the aforementioned helices with a varying harmonic center starting from the orientation quaternions of the initial conformation of the SMD simulation and changing towards those of a target structure that was built based on a homology model of the MFS protein lactose permease, whose crystal structure is in the OF state (PDB: 3O7Q).^66^ The biasing potential was 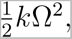 where *k* = 10, 000 *kcal/*(*molÅ*^2^) was the harmonic constant and Ω was the geodesic distance between the current orientation quaternion and a nonlinear interpolation of the initial and target orientation quaternions based on the Riemannian geometry. The outcome of the SMD simulation was equilibrated again for approximately 200 ns to determine the stability of the generated OF state of GkPOT. Prior to this follow-up equilibrium simulation, the final harmonic restraint on orientation quaternions were released gradually within a 15-ns SMD simulation with a fixed center and a variable force constant from 10,000 to 0 *kcal/*(*molÅ*^2^).

## Results and Discussion

Characterizing structural transitions of membrane transporters without compromising the detailed chemical description of these systems and their environments is computationally challenging mainly due to the prohibitively long timescales involved in such processes. Here, we were able to capture the GluT1 IF*→*OF transition using unbiased equilibrium MD for both sets of simulations. However, the sets undergo a conformational change to the OF conformation at different time scales. Within the first 200 ns for Set 1 and within the last 300 ns for Set 2.

On the other hand, a previously reported study of GkPOT^48^ using 400 ns simulations in various *apo* and substrate-bound conditions revealed the shortcomings of unbiased MD for the study of global conformational changes in this protein as the IF*→*OF transition was not captured using such simulations because the time scale of the simulation was much shorter than the associated transition between the IF*→*OF conformational states.^48^ As a result, significant changes in the protein global conformational dynamics were not able to be captured due to a lack of statistically significant fluctuations. ^48^ Here we used the equilibrated structure of *apo* GkPOT^48^ to initiate nonequilibrium pulling SMD simulations to capture the IF*→*OF conformational transition as described in Theoretical Methods. To accomplish this task a number of TM helix biasing protocols were conducted to steer the conformational change to the unknown OF conformation. The combination of H1, H2, H7 and H8 provided the most successful attempt, evaluated by the stability of the resulting OF conformation; i.e., once equilibrated, the resulting conformation from the SMD simulation should stay in an OF conformation with an open periplasmic gate and a closed cytoplasmic gate. Similar to GkPOT, the simulations of GlpT that were previously conducted by Moradi et al.,^6^ used a similar biasing protocol which only involved the TM helices H1 and H7. This protocol was trialed in our GkPOT studies (see Supporting Information); however, the resulting OF conformation was not stable compared to that generated using the H1/H2/H7/H8 protocol. Here, we have also shown GlpT simulations that were previously reported by Moradi et al.^6^for comparison to our simulations conducted on GluT1 and GkPOT.

### Global Protein Conformational Dynamics

Previously reported simulations of GlpT by Moradi et al.,^6^ revealed an important role for TM helices H1, H5, H7, and H11 regarding conformational changes among them by defining interhelical angle pairings between H1 and H7 (⟨H1,H7⟩) and between H5 and H11 (⟨H5,H11⟩). The former is directly involved in periplasmic/extracellular gating and the latter is involved in cytoplasmic/intracellular gating. Figure 2 provides insight into the IF and OF conformations of the respective proteins and helices involved in the cyto-/intra- and periplasmic/extracellular gating with the TM helix pairings colored blue (H1, H7) and red (H5, H11). All proteins studied share a very similar topology (12 TM structure and similar openings at the two gates) despite belonging to a different class of the MFS. GluT1 (Fig. 2A), unlike GkPOT and GlpT, contains an additional domain, the ICH, at the intracellular gate.

**Figure 2:**
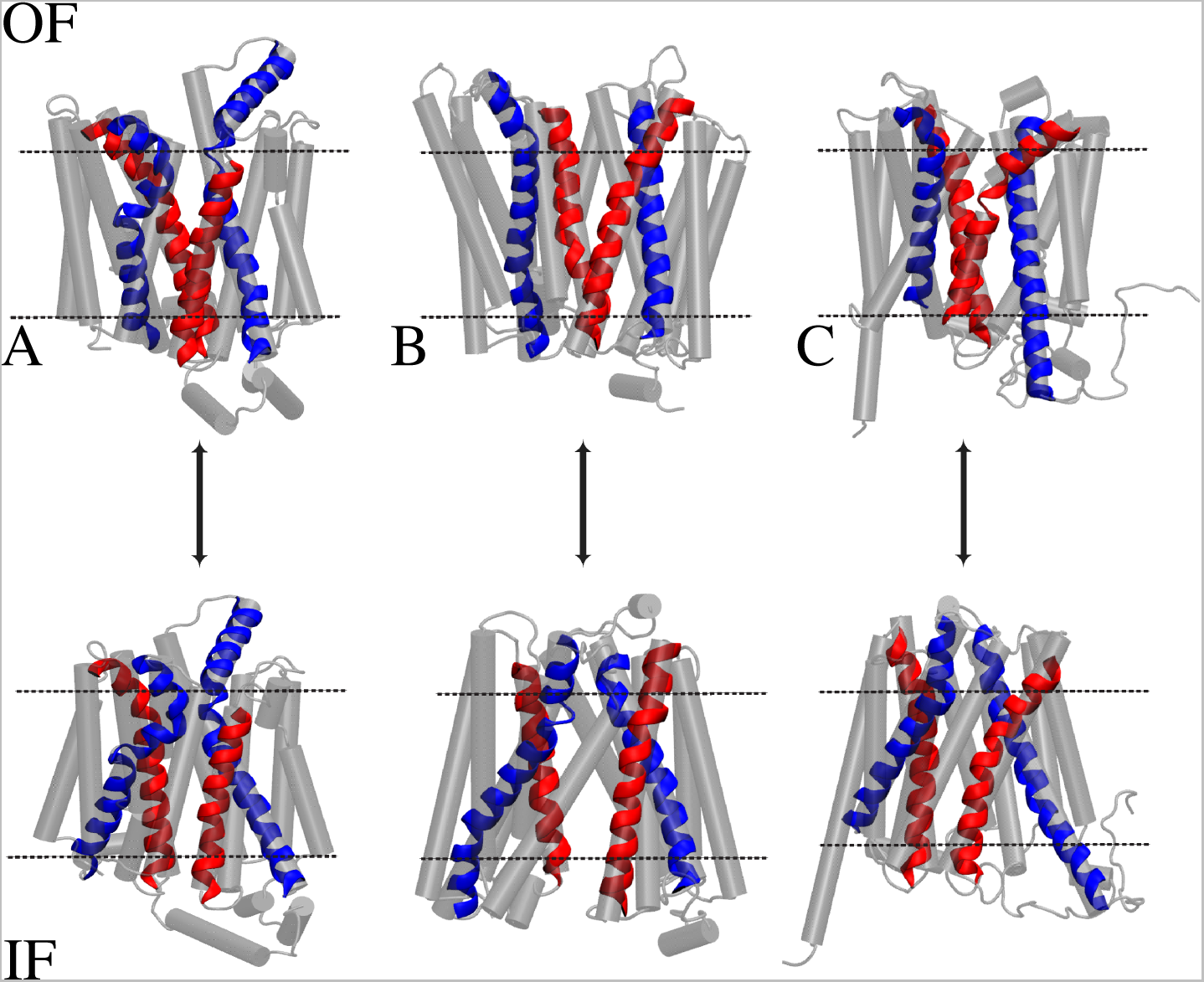
MFS transporters GluT1 (A), GkPOT (B), and GlpT (C) in the OF (top) and IF (bottom) conformations. Helices 1 and 7 are colored in blue and helices 5 and 11 are colored in red. The remaining helices are colored grey to extenuate the changes in the helices that play dominant roles in the cytoplasmic and periplasmic/extracelluar and intracellular gating.

Figure 3 describes the changes associated with the interhelical angles for GluT1, GkPOT, and GlpT. The nature of the three sets of simulations are different; however, one can determine a trend regarding the changes in the interhelical angles in all three sets of simulations that will be discussed below. We have displayed both SMD and the post equilibration data for GkPOT so it can be seen that GkPOT transitions to a stable OF conformation. For GluT1, both Sets 1 and 2 undergo conformational changes at very quick timescales. GluT1 transitions from the IF conformation to an OCC conformation and then to the OF conformation within 100 ns in Set 1 and at *∼*400 ns for Set 2. The transitions are observed by changes in the interhelical angles for ⟨H1,H7⟩ and ⟨H5,H11⟩ as indicated in Figure 3 panels A-B.

**Figure 3:**
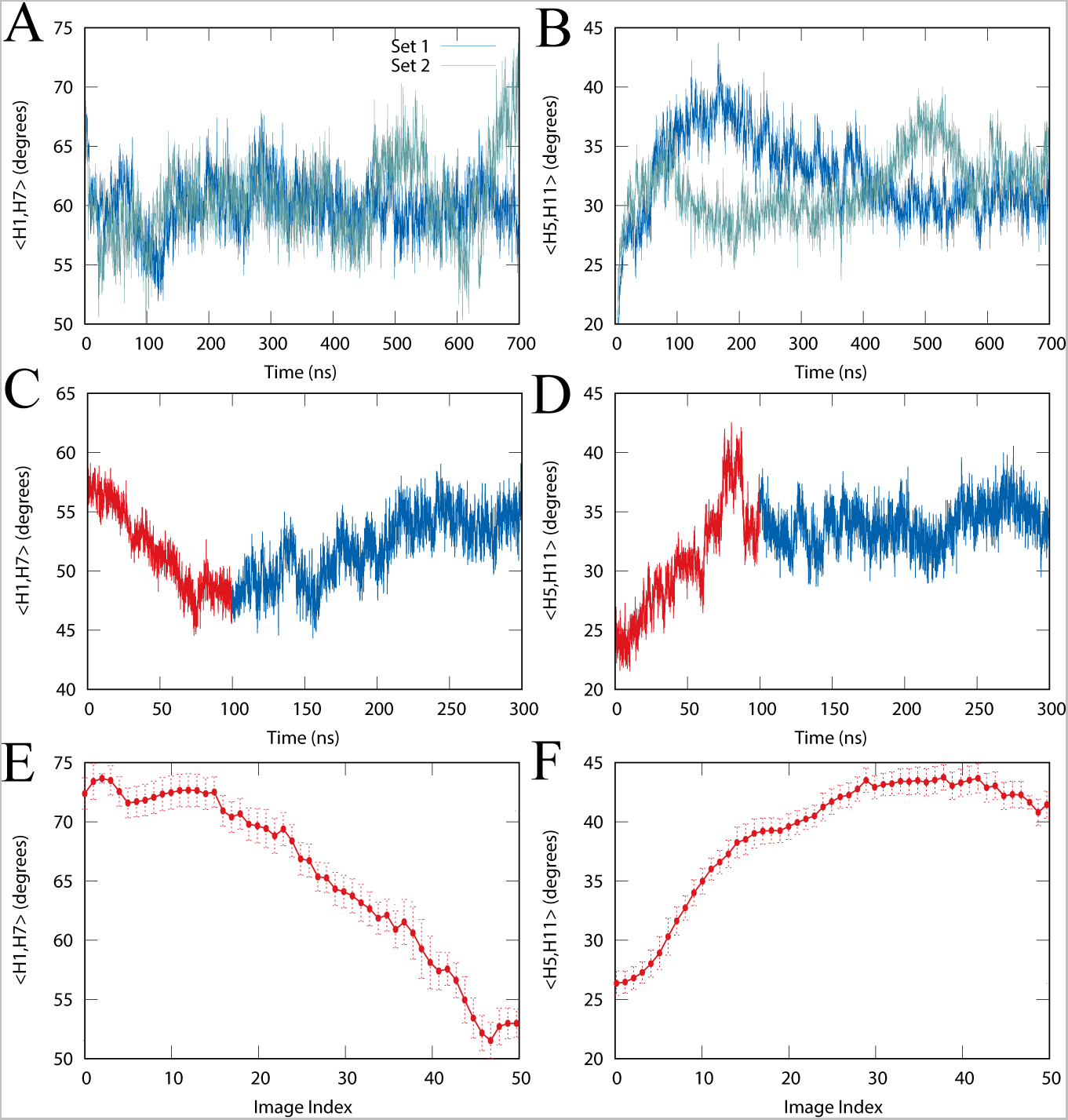
Changes in interhelical angles ⟨H1,H7⟩ and ⟨H5,H11⟩ in GluT1 (A,B), GkPOT (C,D), and GlpT (E,F) simulations. GluT1 time series plots are based on unbiased equilibrium simulations. GkPOT time series include SMD (red) and follow-up release (blue) simulations. GlpT values are based on bias-exchange umbrella sampling simulations, where images 0 and 50 correspond to IF and OF states, respectively.^6^

Our simulations on GluT1 and the simulations conducted by Moradi et al. ^6^ on GlpT show pronounced rotational changes in the interhelical angle between H1 and H7. For set 2 in GluT1, during the conformational transition, ⟨H1,H7⟩ decreases from 68*^◦^* to 52*^◦^* and is followed by a rotational increase to *∼*74*^◦^*. GkPOT undergoes a similar conformational change to that observed in GlpT along the interhelical angles. We note that during the post SMD equilibration the ⟨H1,H7⟩angle increases again to some extent in GkPOT simulations. This is potentially due to the fact that the change imposed on the ⟨H1,H7⟩ angle based on the lactose permease structure (see Theoretical Methods) is more than the actual change in ⟨H1,H7⟩ during the IF-OF transition. The outcome after the equilibration, is more closely similar to that of GluT1, which is less than 10 degrees. On the other hand, the ⟨H1,H7⟩ change in GlpT is about 20 degrees. For ⟨H5,H11⟩, the GluT1 transporter shows the highest change. However, all three proteins show a significant correlation between the changes in ⟨H1,H7⟩ and ⟨H5,H11⟩, in line with the rocker-switch mechanism.

### Local Conformational Changes

The GluT1 extracelluar gating is directly influenced by the interactions between TM helices H1 and H7. This is consistent with both GkPOT and GlpT (Fig. 3). However, unlike GkPOT and GlpT, the interactions between H1 and H7 take place on the extracellular side of the protein as opposed to being in the TM region (Fig. 4A). The salt bridges observed in the TM region of both GkPOT and GlpT are necessary for protein stability and play a key role in substrate translocation (Fig. 4).

**Figure 4:**
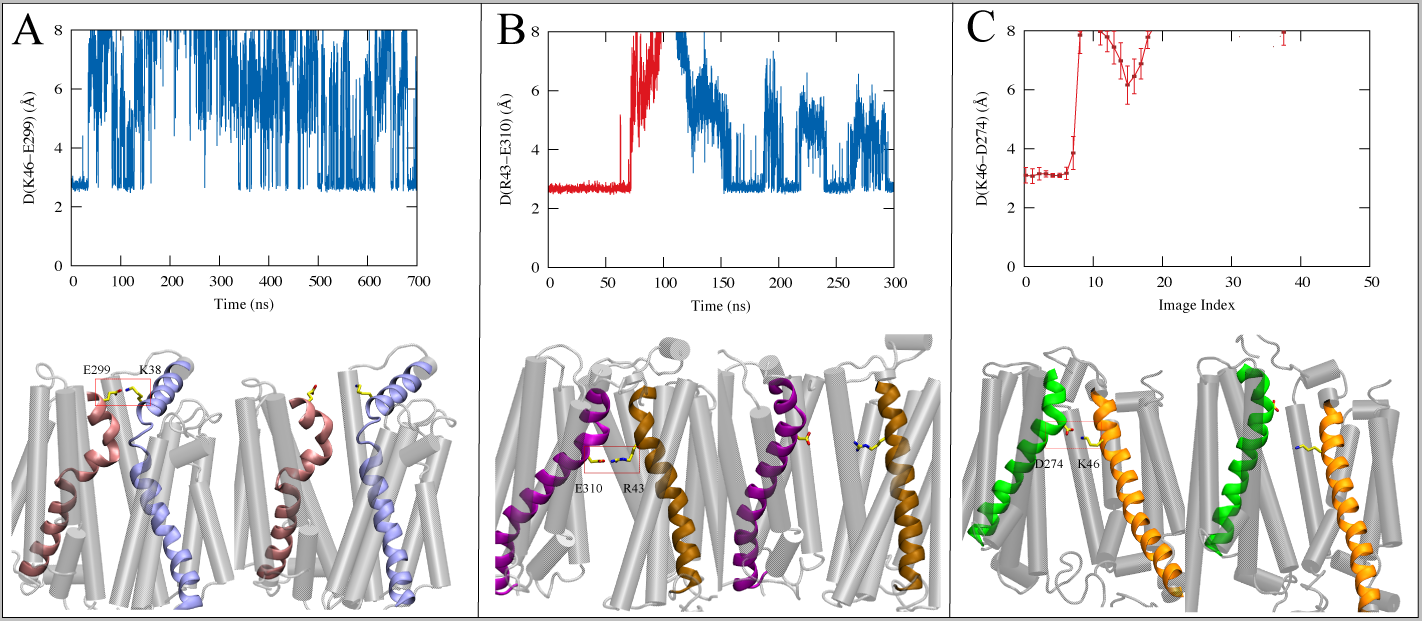
Minimum donor-acceptor distance measurements between the key inter-bundle salt-bridge forming amino acids (A) K38 and E299 in GluT1 (Set 1), (B) R43 and E310 in GkPOT, and (C) K46 and D274 in GlpT.

A salt bridge between Lys 38 in H1 and Glu 299 in H7 is observed in the very beginning of the GluT1 simulation (Fig. 4A and Supplementary Information), as the protein remains in the IF conformation. After moving to the occluded conformation it can be seen that the salt bridge tries to reform. Additional salt-bridge forming residues have also been observed and appear to be directly involved in allowing GluT1 to move into the OF conformation. For GluT1, the ICH provides the cytoplasmic gating by moving into a position to interact with both H5 and H11. The formation of two very strong salt bridges with H5 and H11 (Glu 243 and Arg 153, Glu 247 and Arg 400) (Supplementary Information), allow for the ICH to remain locked into position. From this stability we observe the interhelical angles between H5-H11 to exhibit the same angular change of about 15-20 degrees during the transitions between IF and OF conformations.

As GkPOT transitions toward the OF state, an important inter-bundle salt bridge between R43 in H1 and E310 in H7 becomes disrupted, and appears to break after 75 ns of the SMD pulling (Fig. 4B). The salt bridge forms again after the protein reaches the equilibrated OF conformation, although the strength of this salt bridge is extremely weakened. The salt bridge forms and breaks multiple times during the equilibrated portion of the simulation.

GlpT, on the other hand, has the strongest correlation between the local events (i.e., salt bridge formation/disruption) and global conformational changes (interhelical angle changes). The interbundle salt bridges seem to play an important role in the transport mechanism of MFS transporters; however, the exact location and the exact function of these salt bridges make it quite difficult to draw any conclusion on this issue. The interbundle salt bridges seem to stabilize the IF conformation in all three proteins and the IF*→*OF transition seems to require the disruption of this salt bridge. The OF state, however, may or may not be associated with the presence or absence of the salt bridge.

### Water Accessibility at Cytoplasmic and Periplasmic Gates

Figure 5 depicts the change in water count along the protein pore and the average water count at the cytoplasmic and periplasmic gates of GluT1 during different stages of the equilibrium simulations. It can be seen that GluT1 transitions very quickly to an OCC conformation, within the first few nanseconds of the simulations, as indicated in the water profiles of Figure 5 (A-B). Water profiles from different stages of the equilibrium simulations show an increase in water found at the extracellular gate which is indicated in panels C-D. The average water counts at the intra- and extracellular gates are also shown for the full 700 ns time series and at different time intervals after the GluT1 has transitioned to the OF conformation (Fig. 5E-H). From this, it is clear that there is a separation in the amount of water present at the extracellular gate in both simulation sets shown in Figure 5 (E-H). Additionally, the extracellular gate appears to close briefly for Set 2 (Fig. 5H) but reverts back as the protein is trying to stabilize in the OF conformation at time 550 ns.

**Figure 5:**
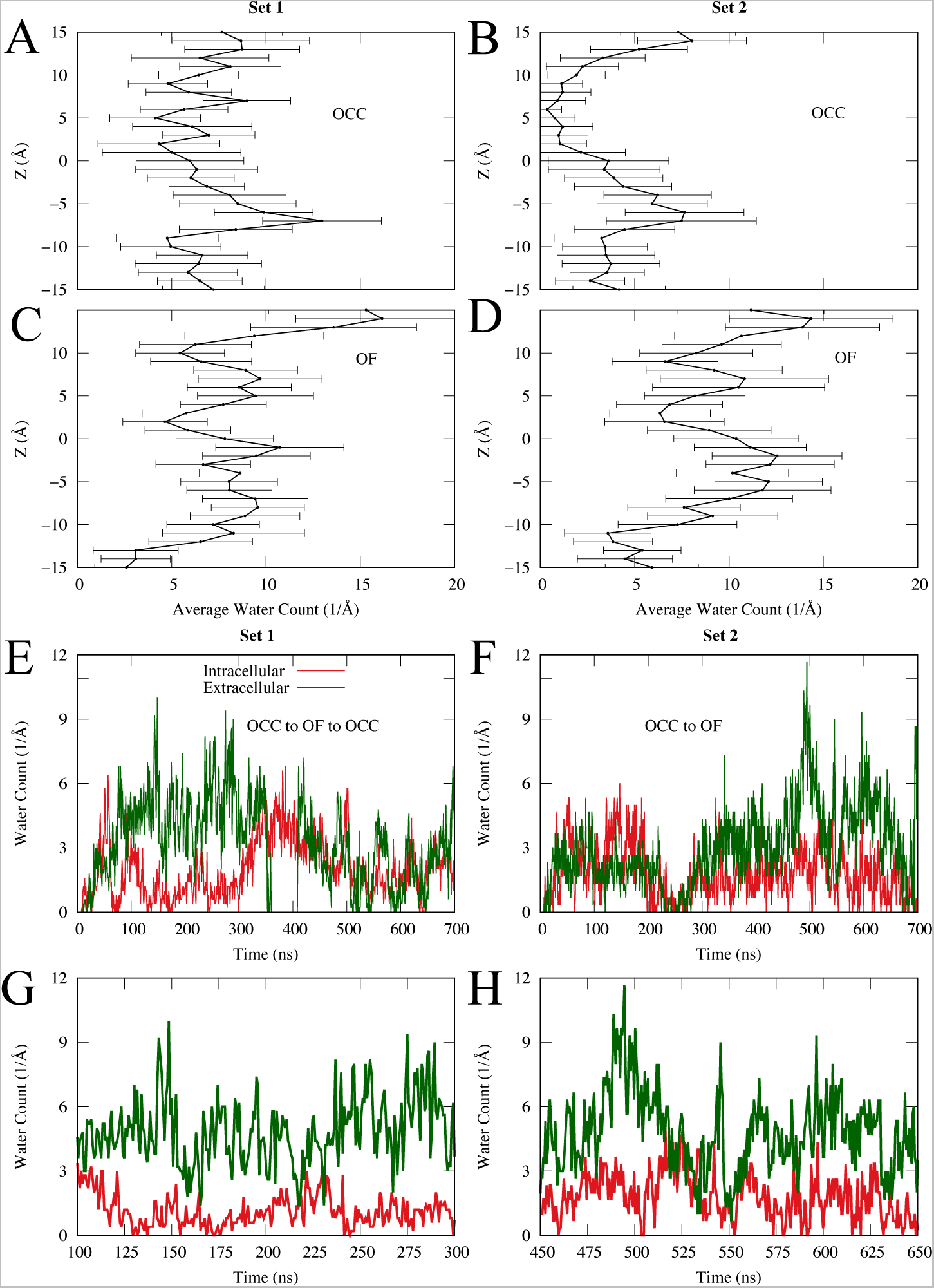
GluT1 water accessibility of the two simulation sets along the pore (A-D) and at the extra-(green) and intracellular (red) gates during different stages of the equilibrium simulations (E-H): (A, B) 100 ns of equilibrium simulation of GluT1 in the OCC state prior to transitioning to OF conformation. (C, D) Average water count over 20 ns of simulation after transitioning to the OF conformation. (E, F) Time series plots showing the water count at the extra-(green) and intracellular (red) gates for both sets for the entire simulation time. (G, H) 200 ns of the OF water count at the associated time intervals for a given set.

Similarly, Figure 6 depicts the change in water count along the protein pore and the average water count at the cytoplasmic and periplasmic gates of GkPOT from different stages of the equilibrium and nonequilibrum simulations. Figure 6 panels A-B show the comparison of the water along the protein pore when in the IF and OF conformations both before and after the nonequilibrium simulation. The comparison between the two states shows a much broader degree of the water at the cytoplasmic side of the protein which is attributed to GkPOT remaining in the IF state the entirety of the unbiased simulation. On the other hand, the OF conformation from the post unbiased simulation shows a water profile that has water both at the cytoplasmic and periplasmic sides. We attribute this profile to the overall function of GkPOT and the ability to accommodate different substrates in different binding orientations which would account for a slightly “leaky” bottleneck at both gates. Additionally, it can be seen that the average water along the pore in the GkPOT OF conformation is very comparable to that of the OF conformation of GluT1. Our observations found from panels A-B are further advanced by panels C-E which show water accessibility at the different stages of the simulations. The average water count at the periplasmic gate trends to be above the average water count found at the cytoplasmic gate during the nonequilibrium simulation and continues to remain higher during the post nonequilibrium simulation. Admittedly, panel E only represents the first 100ns of the simulation, however, panel B shows the water profile for all 200 ns of the unbiased simulation.

**Figure 6:**
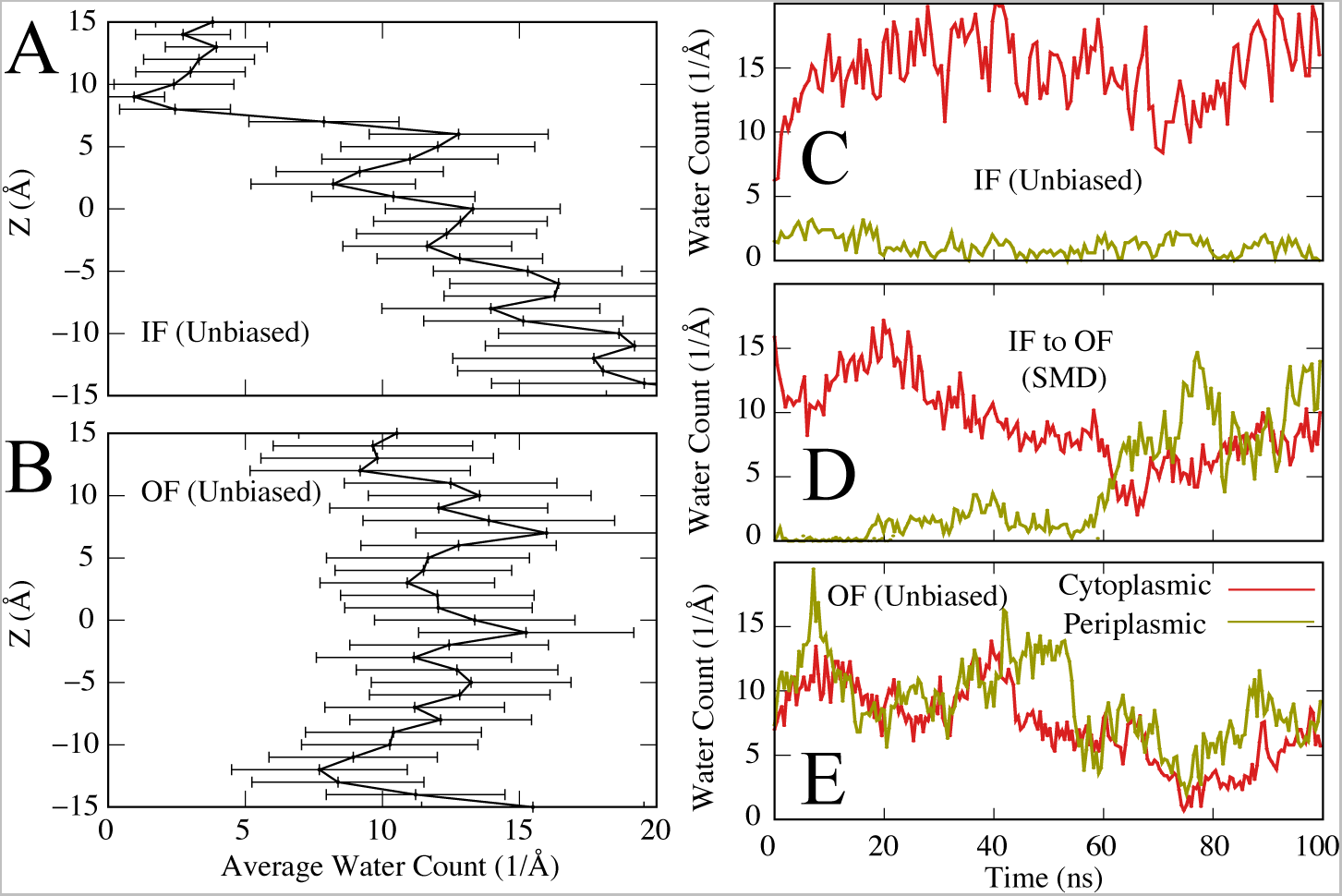
GkPOT water accessibility along the pore (A,B), at peri-(bronze) and cytoplasmic (red) gates during different stages of equilibrium and nonequilibrium simulations: (C) 100 ns of equilibrium simulation of GkPOT in the IF state prior to SMD (D) 100 ns of SMD as described previously for GkPOT, and (E) the first 100 ns of follow-up unbiased MD after transitioning to the OF state. The water profiles and the error bars in (A) and (B) are generated from the same simulations used for panels (C) and (E).

The water profiles and time series of the water counts at both the peri-/extra- and cytoplasmic/intracellular gates further supports the alternating-access mechanism in both GluT1 and GkPOT. Although we do not observe GluT1 transition back towards the IF conformation in the two sets of simulations, the trends indicated in Figures 5 and 6 show that the proteins are not open to both sides of the membrane at the same time and suggests that the conformational changes of the *apo* proteins follow ideal gating modes.

## Conclusion

Here, we have performed all-atom equilibrium MD simulations of GluT1 and all-atom nonequilibrium MD simulations of GkPOT as well as compared our simulations to previously reported simulations of GlpT performed by Moradi et al.^6^ Through this study we have demonstrated a thorough comparison of the conformational dynamics of the three different classes of MFS transporters; i.e., symporters, uniporters, and antiporters, with emphasis placed on the simulation of the *apo* protein.

The three systems (classes) of MFS transporters examined are quite different in terms of the timescale of the IF-OF transition; however, we observe various similarities between the three systems including the involvement of specific TM helices that undergo rotational changes along the transition pathways. These include, the H1/H7 helices which are involved in periplasmic/extracellular gating and the H5/H11 helices, involved in cytoplasmic/intracellular gating. Additionally, these three systems contain an interbundle salt bridge that forms between the H1 and H7. The formation of this interbundle salt bridge stabilizes these three systems of MFS transporters in the IF state. Admittedly, there are notable differences between these three systems including: (1) the smaller change of ⟨H1,H7⟩angle in GkPOT as compared to GluT1 and GlpT, (2) the larger change of ⟨H5,H11⟩ angle in GluT1 as compared to GkPOT and GlpT, and (3) the formation of the interbundle salt bridge in the OF state for the GkPOT protein as compared to GluT1 and GlpT.

The presented study provides a detailed picture of the similarities and differences associated with the IF*↔*OF conformational transition of three transporters from three distinct classes of MFS superfamily. However, additional research needs to be conducted involving the transporters with their substrate(s) bound to gain a better understanding of the transition between the IF*↔*OF conformational states.

## Acknowledgement

This research is supported by the University of Arkansas, Fayetteville. This research is part of the Blue Waters sustained-petascale computing project, which is supported by the National Science Foundation (awards OCI-0725070 and ACI-1238993) and the state of Illinois. Blue Waters is a joint effort of the University of Illinois at Urbana-Champaign and its National Center for Supercomputing Applications. This work also used the Extreme Science and Engineering Discovery Environment (allocation MCB150129), which is supported by National Science Foundation grant number ACI-1548562. This research is also supported by the Arkansas High Performance Computing Center which is funded through multiple National Science Foundation grants and the Arkansas Economic Development Commission.

## References

(1) Reddy, V. S.; Shlykov, M. A.; Castillo, R.; Sun, E. I.; Saier, M. H. The major facilitator superfamily (MFS) revisited. FEBS J. 2012, 279, 2022–2035.

(2) Pao, S. S.; Paulsen, I. T.; Saier, M. H. Major facilitator superfamily. Microbiol. Mol. Biol. Rev. 1998, 62, 1 LP–34.

(3) Law, C. J.; Almqvist, J.; Bernstein, A.; Goetz, R. M.; Huang, Y.; Soudant, C.; Laaksonen, A.; Hovmöller, S.; Wang, D.-N. Salt-bridge dynamics control substrate-induced conformational change in the membrane transporter GlpT. J. Mol. Biol. 2008, 378, 828–839.

(4) Huang, M., Y. Lemieux; Song, J.; Auer, M.; Wang, D. Structure and mechanism of the glycerol-3-phosphate transporter from Escherichia coli. Science 2003, 301, 616–620.

(5) Drew, D.; North, R.; Nagarathinum, K.; Tanabe, M. Structures and general transport mechanisms by the major facilitator superfamily (MFS). Chem. Rev. 2021, 121, 5289– 5335.

(6) Moradi, M.; Enkavi, G.; Tajkhorshid, E. Atomic-level characterization of transport cycle thermodynamics in the glycerol-3-phosphate:phosphate antiporter. Nat. Commun. 2015, 6, 8393.

(7) Madej, M. G. Function, structure, and evolution of the major facilitator superfamily: the LacY manifesto. Advances in Biology 2014, 523591.

(8) Feng, J.; Selvam, B.; Shukla, D. How do antiporters exchange substrates across the cell membrane? An atomic-level description of the complete exchange cycle in NarK. Structure 2021, 29, 922–933.

(9) Selvam, B.; Mittal, S.; Shukla, D. Free energy landscape of the complete transport cycle in a key bacterial transporter. ACS Cent. Sci. 2018, 4, 1146–1154.

(10) Lee, J.; Sands, Z. A.; Biggin, P. C. A numbering system for MFS transporter proteins. Front. Mol. Biosci. 2016, 3.

(11) Abramson, J.; Smirnova, I.; Kasho, V.; Verner, G.; Kaback, H. R.; Iwata, S. Structure and mechanism of the lactose permease of Escherichia coli. Science 2003, 301, 610–615.

(12) Guan, L.; Hu, Y.; Kaback, R. Aromatic stacking in the sugar binding site of the lactose permease. Biochemistry 2003, 42, 1377–1382.

(13) Guan, L.; Kaback, R. Lesssons from lactose permease. Annu. Rev. Biophys. Biomol. Struct. 2006, 35, 67–91.

(14) Stelzl, L. S.; Fowler, P. W.; Sansom, M. S.; Beckstein, O. Flexible gates generate occluded intermediates in the transport cycle of LacY. J. Mol. Biol. 2014, 426, 735– 751.

(15) Lawrence, M. G.; Lindahl, L.; Zengel, J. M. Effects on translation pausing of alterations in protein and RNA components of the ribosome exit tunnel. J. Bacteriol. 2008, 190, 5862–5869.

(16) Quistgaard, E. M.; Löw, C.; Moberg, P.; Tŕesaugues, L.; Nordlund, P. Structural basis for substrate transport in GLUT-homology family of monosaccharide transporters. Nat. Struct. Mol. Biol. 2013, 20, 766–768.

(17) Yin, Y.; Jensen, M.; Tajkhorshid, E.; Schulten, K. Sugar binding and protein conformational changes in lactose permease. Biophys. J. 2006, 91, 3972–3985.

(18) Yan, N. Structural advances for the major facilitator superfamily (MFS) transporters. Trends Biochem. Sci. 2013, 38, 151–159.

(19) Lemieux, J. Eukaryotic major facilitator superfamily transporter modeling based on the prokaryotic GlpT crystal structure (Review). Mol. Membr. Biol. 2007, 24, 333–341.

(20) Selvam, B.; Feng, J.; Shukla, D. Atomistic insights into the mechanism of dual affinity switching in plant nitrate transporter NRT1.1. BioRxiv 2022,

(21) Wood, I. S.; Trayhurn, P. Glucose transporters (GLUT and SGLT): expanded families of sugar transport proteins. BR J. Nutr. 2003, 89, 3–9.

(22) Patlak, C. Contributions to the theory of active transport: II. The gate type non-carrier mechanism and generalizations concerning tracer flow, efficiency, and measurement of energy expenditure. Bull. Math. Biol. 1957, 19, 209–235.

(23) Vidaver, G. Inhibition of parallel flux and augmentation of counter flux shown by transport models not involving a mobile carrier. J. Theor. Biol. 1966, 10, 301–306.

(24) Jardetzky, O. Simple allosteric model for membrane pumps. Nature 1966, 211, 969–970.

(25) Ryan, R.; Vandenberg, R. Elevating the alternating-access model. Nat. Struct. Mol. Biol. 2016, 23, 187–189.

(26) Chan, M.; Shukla, D. Markov state modeling of membrane transport proteins. J. Struct. Biol. 2021, 213, 107800.

(27) Beckstein, O.; Naughton, F. General principles of secondary active transporter function. Biophys. Rev. 2021, 3, 011307.

(28) Weyand, S.; Shimamura, T.; Beckstein, O.; Sansom, M.; Iwata, S.; Henderson, P.; Cameron, A. The alternating access mechanism of transport as observed in the sodium-hydantoin transporter Mhp1. J. Synchrotron Radiat. 2011, 18, 20–23.

(29) Sauve, S.; Williamson, J.; Polasa, A.; Moradi, M. Ins and outs of rocker switch mechanism in major facilitator superfamily of transporters. Membranes 2023, 13, 462.

(30) Yin, Y.; He, X.; Szewczyk, P.; Nguyen, T.; Chang, G. Structure of the multidrug transporter EmrD from Escherichia Coli. Science 2006, 312, 741–744.

(31) Yan, H.; Huang, W.; Yan, C.; Gong, X.; Jiang, S.; Zhao, Y.; Wang, J.; Shi, Y. Structure and mechanism of a nitrate transporter. Cell Rep. 2013, 3, 716–723.

(32) Newstead, S. Towards a structural understanding of drug and peptide transport within the proton-dependent oligopeptide transporter (POT) family. Biochemical Society Transactions 2011, 39, 1353 LP – 1358.

(33) Solcan, N.; Kwok, J.; Fowler, P. W.; Cameron, A. D.; Drew, D.; Iwata, S.; Newstead, S. Alternating access mechanism in the POT family of oligopeptide transporters. EMBO J. 2012, 31, 3411–3421.

(34) Guettou, F.; Quistgaard, E. M.; Tresaugues, L.; Moberg, P.; Jorgerschold, C.; Jong, A. J.; Nordlund, P.; Loew, C. Structural insights into substrate recognition in proton-dependent oligopeptide transporters. EMBO Rep. 2013, 14, 804–810.

(35) Thorens, B.; Mueckler, M. Glucose transporters in the 21st Century. Am. J. Physiol. – Endoc. M. 2010, 298, E141–E145.

(36) Deng, D.; Xu, C.; Sun, P.; Wu, J.; Yan, C.; Hu, M.; Yan, N. Crystal structure of the human glucose transporter GLUT1. Nature 2014, 510, 121–125.

(37) Deng, D.; Sun, P.; Yan, C.; Ke, M.; Jiang, X.; Xiong, L.; Ren, W.; Hirata, K.; Yamamoto, M.; Fan, S. et al. Molecular basis of ligand recognition and transport by glucose transporters. Nature 2015, 526, 391.

(38) Yan, N. A glimpse of membrane transport through structures—advances in the structural biology of the GLUT glucose transporters. J. Mol. Biol. 2017, 429, 2710–2725.

(39) Yan, N. Structural advances for the major facilitator superfamily (MFS) transporters. Trends Biochem. Sci. 2013, 38, 151–159.

(40) Doki, S.; Kato, H. E.; Solcan, N.; Iwaki, M.; Koyama, M.; Hattori, M.; Iwase, N.; Tsukazaki, T.; Sugita, Y.; Kandori, H. et al. Structural basis for dynamic mechanism of proton-coupled symport by the peptide transporter POT. Proc. Natl. Acad. Sci. USA 2013, 110, 11343–11348.

(41) Lichtinger, S. M.; Parker, J. L.; Newstead, S.; Biggin, P. C. The mechanism of mammalian proton-coupled peptide transporters. BioRxiv 2024.,

(42) Yang, Q.; Ma, Y.; Zhao, Y.; She, Z.; Wang, L.; Li, J.; Wang, C.; Deng, Y. Accelerated drug release and clearance of PEGylated epirubicin liposomes following repeated injections: a new challenge for sequential low-dose chemotherapy. Int. J. Nanomedicine 2013, 8, 1257–1268.

(43) Lyons, J. A.; Parker, J. L.; Solcan, N.; Brinth, A.; Li, D.; Shah, S. T.; Caffrey, M.; Newstead, S. Structural basis for polyspecificity in the POT family of proton-coupled oligopeptide transporters. EMBO Rep. 2014, 15, 886 – 893.

(44) Newstead, S.; Drew, D.; Cameron, A. D.; Postis, V. L.; Xia, X.; Fowler, P. W.; Ingram, J. C.; Carpenter, E. P.; Sansom, M. S.; McPherson, M. J. et al. Crystal structure of a prokaryotic homologue of the mammalian oligopeptide-proton symporters, PepT1 and PepT2. EMBO J. 2011, 30, 417–426.

(45) Quistgaard, E. M.; Martinez Molledo, M.; Low, C. Structure determination of a major facilitator peptide transporter: Inward facing PepTSt from Streptococcus thermophilus crystallized in space group P3121. PLOS ONE 2017, 12, 1–20.

(46) Elvin, C. M.; Hardy, C. M.; Rosenberg, H. Pi exchange mediated by the GlpT-dependent sn-glycerol-3-phosphate transport system in Escherichia coli. J. Bacteriol. 1985, 161, 1054– 1058.

(47) Law, C. J.; Almqvist, J.; Bernstein, A.; Goetz, R. M.; Huang, Y.; Soudant, C.; Laaksonen, A.; Hovmöller, S.; Wang, D.-N. Salt-bridge dynamics control substrate-induced conformational change in the membrane transporter GlpT. J. Mol. Biol. 2008, 378, 828–839.

(48) Immadisetty, K.; Hettige, J.; Moradi, M. What can and cannot be learned from molecular dynamics simulations of bacterial proton-coupled oligopeptide transporter GkPOT? J. Phys. Chem. B 2017, 121, 3644–3656.

(49) Immadisetty, K.; Moradi, M. Mechanistic picture for chemomechanical coupling in a bacterial proton-coupled oligopeptide transporter from Streptococcus Thermophilus. J. Phys. Chem. B 2021, 125, 9738–9750.

(50) Galochkina, T.; Ng Fuk Chong, M.; Challali, L.; Abbar, S.; Etchebest, C. New insights into GluT1 mechanics during glucose transfer. Sci. Rep. 2019, 9, 998.

(51) Isralewitz, B.; Stone, J.; Schulten, K. Timeline: an interactive raster plot to identify events in molecular dynamics trajectories. 2011, Submitted.

(52) Schlick, T.; Li, B.; Olson, W. The influence of salt on the structure and energetics of supercoiled DNA. Biophys. J. 1994, 67, 2146–2166.

(53) Moradi, M.; Tajkhorshid, E. Mechanistic picture for conformational transition of a membrane transporter at atomic resolution. Proc. Natl. Acad. Sci. USA 2013, 110, 18916–18921.

(54) Moradi, M.; Tajkhorshid, E. Computational recipe for efficient description of large-scale conformational changes in biomolecular systems. J. Chem. Theory Comput. 2014, 10, 2866–2880.

(55) Fakharzadeh, A.; Moradi, M. Effective riemannian diffusion model for conformational dynamics of biomolecular systems. J. Phys. Chem. Lett. 2016, 7, 4980–4987.

(56) Jo, S.; Kim, T.; Im, W. Automated builder and database of protein/membrane complexes for molecular dynamics simulations. PLoS One 2007, 2, e880.

(57) Lee, J.; Cheng, X.; Swails, J. M.; Yeom, M. S.; Eastman, P. K.; Lemkul, J. A.; Wei, S.; Buckner, J.; Jeong, J. C.; Qi, Y. et al. CHARMM-GUI input generator for NAMD, GROMACS, AMBER, OpenMM, and CHARMM/OpenMM simulations using the CHARMM36 additive force field. J. Chem. Theory Comp. 2016, 12, 405–413.

(58) Jorgensen, W. L.; Chandrasekhar, J.; Madura, J. D.; Impey, R. W.; Klein, M. L. Comparison of simple potential functions for simulating liquid water. J. Chem. Phys. 1983, 79, 926–935.

(59) Klauda, J. B.; Venable, R. M.; Freites, J. A.; O’Connor, J. W.; Tobias, D. J.; Mondragon-Ramirez, C.; Vorobyov, I.; MacKerell Jr., A. D.; Pastor, R. W. Update of the CHARMM all-atom additive force field for lipids: Validation on six lipid types. J. Phys. Chem. B 2010, 114, 7830–7843.

(60) Best, R. B.; Zhu, X.; Shim, J.; Lopes, P. E. M.; Mittal, J.; Feig, M.; MacKerell, A. D. Optimization of the additive CHARMM all-atom protein force field targeting improved sampling of the backbone *ϕ*, ψ and side-chain χ1 and *χ*2 dihedral angles. J. Chem. Theory Comp. 2012, 8, 3257–3273.

(61) Phillips, J. C.; Braun, R.; Wang, W.; Gumbart, J.; Tajkhorshid, E.; Villa, E.; Chipot, C.; Skeel, R. D.; Kale, L.; Schulten, K. Scalable molecular dynamics with NAMD. J. Comp. Chem. 2005, 26, 1781–1802.

(62) Reid, J. K. In Large Sparse Sets of Linear Equations; Reid, J. K., Ed.; Academic Press: London, 1971; pp 231–254.

(63) Martyna, G. J.; Tobias, D. J.; Klein, M. L. Constant pressure molecular dynamics algorithms. J. Chem. Phys. 1994, 101, 4177–4189.

(64) Feller, S. E.; Zhang, Y.; Pastor, R. W.; Brooks, B. R. Constant pressure molecular dynamics simulation: The Langevin piston method. J. Chem. Phys. 1995, 103, 4613– 4621.

(65) Darden, T.; York, D.; Pedersen, L. G. Particle mesh Ewald: An *N ·*log(*N*) method for Ewald sums in large systems. J. Chem. Phys. 1993, 98, 10089–10092.

(66) Dang, S.; Sun, L.; Huang, Y.; Lu, F.; Liu, Y.; Gong, H.; Wang, J.; Yan, N. Structure of a fucose transporter in an outward-open conformation. Nature 2010, 467, 734–738.

