## Supporting Information for "Conformational Transition Pathways in Major Facilitator Superfamily Transporters"

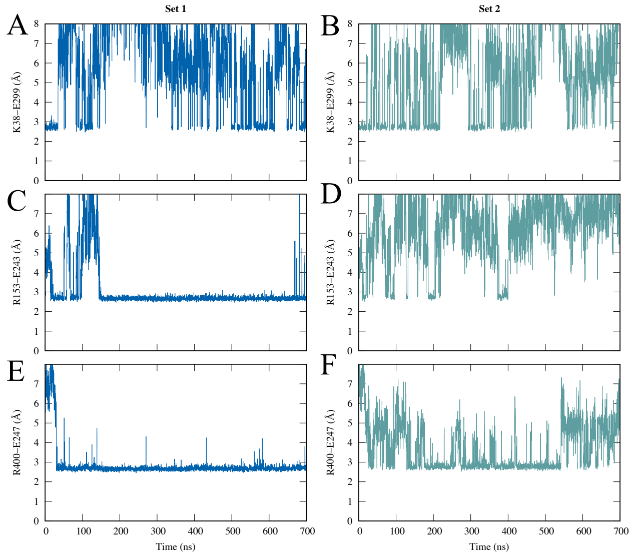


Figure S1: GluT1 minimum donor-acceptor distance measurements between key inter-bundle salt-bridge forming amino acids in both simulation sets. (A-B) K38 and E299. (B-C) R153 and E243. (E-F) R400 and E247.

### Nonequilibrium Pulling (NEP) Simulations

#### Optimizing the NEP protocol on UP-E310:Apo GKPOT

We considered N-terminal (helices 1-6) and C-terminal (helices 7-12) regions of POTs for the NEP simulations and ignored HA (residues 225-250) and HB (residues 258-280). For the NEP simulations involving the unprotonated-E310:apo form of GKPOT we have used the 140ns equilibrated structure as the starting structure for NEP simulations.

Table S1: Various NEP protocols tested on the UP-E310:***Apo*** form of GKPOT

| Index | Force Constant(K) | Simu. Time(ns) | Work (kcal/mol) |
|---|---|---|---|
| **1,7** | 10,000 | 5 | 200 |
| | 10,000 | 20 | 142 |
| | 10,000 | 100 | |
| | 5000 | 20 | 120 |
| | 4000 | 20 | 115 |
| | 3000 | 20 | 100 |
| | 2000 | 20 | 72 |
| | 1000 | 20 | 53 |
| | 500 | 20 | 28 |
| **1,8** | 10,000 | 20 | 175 |
| **1,9** | 10,000 | 20 | 14 |
| **1,11** | 10,000 | 20 | 190 |
| **1,12** | 10000 | 20 | 170 |
| **2,8** | 10,000 | 5 | 230 |
| **2,8** | 10,000 | 20 | 205 |
| **3,9** | 10,000 | 5 | 13 |
| **3,9** | 10,000 | 20 | 6 |
| **5,11** | 10,000 | 20 | 180 |
| **5,11** | 10,000 | 5 | 225 |
| **6,12** | 10,000 | 5 | 240 |
| **6,12** | 10,000 | 20 | 160 |
| **7,2** | 10,000 | 20 | 190 |
| **7,3** | 10,000 | 20 | 140 |
| **7,4** | 10,000 | 20 | 140 |
| **7,5** | 10,000 | 20 | 132 |
| **7,6** | 10,000 | 20 | 155 |
| | 10,000 | 100 | |
| **4,5,11** | 10,000 | 20 | 215 |
| **1,7,2,8** | 10,000 | 20 | 300 |
| | 10,000 | 100 | |
| | 5000 | 20 | 230 |
| | 5000 | 100 | |
| | 4000 | 20 | 250 |
| | 3000 | 20 | 225 |
| | 2000 | 20 | 180 |
| | 1000 | 20 | 120 |
| **1,7,3,9** | 10,000 | 20 | 180 |
| **1,7,5,11** | 10,000 | 20 | 225 |
| **3,6,9,12** | 10,000 | 20 | 180 |

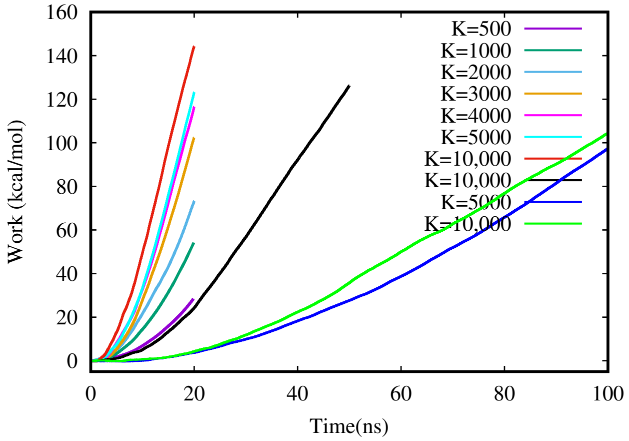


Figure S2: **Optimizing the 1,7 colvars on UP:*Apo* GkPOT.**

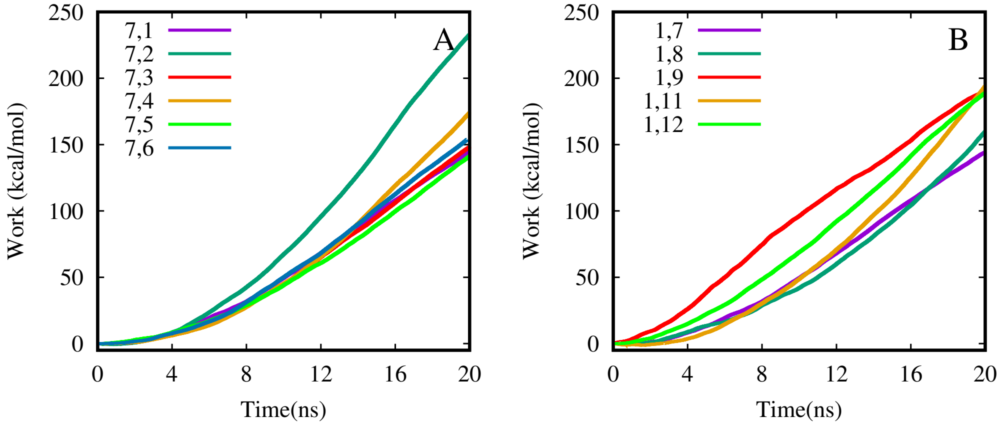


Figure S3: **Comparing the helix-1 and helix-7 colvars combinations on UP:*Apo* GkPOT.**

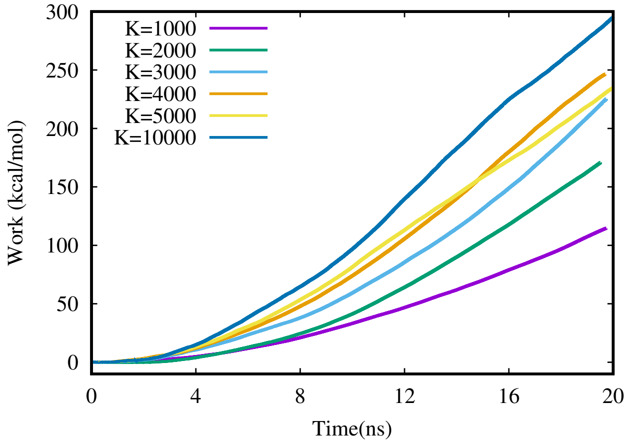


Figure S4: **Application of 1,7,2,8 colvars on UP:*Apo* with different force constants.**
